# Resource-combining costs of being a diet generalist in the super-generalist protist predator *Dictyostelium discoideum*

**DOI:** 10.1101/2023.08.05.552129

**Authors:** P. M. Shreenidhi, Debra A. Brock, Rachel I. McCabe, Joan E. Strassmann, David C. Queller

## Abstract

Consumers lie on a continuum between diet specialization on few resources to being generalist feeders on many resources. Generalism has the clear advantage of having more resources to exploit, but the costs that limit generalism are less clear. We explore two understudied costs of generalism in a super-generalist amoeba predator, *Dictyostelium discoideum*, feeding on naturally co-occurring bacterial prey. Both involve costs of combining different prey. First, amoebas exhibit a reduction in growth rate when they switch from one species of prey bacteria to another, something we call resource-switching costs. These switching costs typically disappear within a day, indicating adjustment to new prey bacteria. Second, amoebas usually divide more slowly on mixtures of bacteria compared to on single bacteria, something we call resource-mixing costs. Both results support that idea that, although amoebas can consume a huge variety of prey, they must use partially different methods and thus must pay costs to handle multiple prey, either sequentially or simultaneously.

**Significance Statement:** Perhaps the most fundamental conflict in nature occurs when one organism consumes another. Diet generalists benefit from the advantage of eating many prey but then must deal with many prey defences. We explore costs associated with a broad diet in a protist microbial predator, *Dictyostelium discoideum*. These predators of bacteria show a delay in growth when switched from one bacteria to another, supporting the hypothesis that they must deploy different strategies. They also experience costs when grown on many bacteria at once, suggesting that the alternative strategies for consuming different prey are partly incompatible with each other. Our findings shed light on the nature of diet generalism and highlight the complexity of predation in the microbial world.

## Introduction

Consumers vary widely in their diet breadth. Some are diet specialists that eat one or few resources, such as koalas that feed only on eucalyptus leaves (1) or snail kites that exclusively hunt apple snails (2). Some are diet generalists that consume many different resources, such as coyotes that feed on many small mammals (3) or spiders that feed on many species of arthropods (4, 5). This variation in diet breadth has important ecological and evolutionary consequences that impact the structure and stability of food-webs (6), the stability of communities to perturbations (7), within- and between-species competition (8), the strength of coevolutionary dynamics (9) and speciation (10).

The obvious benefit to being a diet generalist is the ability to exploit diverse resources, especially when resources are scarce and fluctuate in their availability (11). But there must be associated costs with diet generalism or else all consumers would be generalists. One classic explanation for diet specialization is that “the jack of all-trades is the master of none.” We expect generalists to suffer from trade-offs in resource utilization because of antagonistic pleiotropy, where mutations that are good for exploiting one resource maybe bad on another (12–14). Such trade-offs can also occur via mutation accumulation, where mutations that are neutral on one resource may be detrimental on new resources (15). Generally, there has been surprisingly little support for trade-offs in performance on different resources (12, 16, 17). Other, less studied, costs include that generalists may also be slower to adapt to a given resource because they spend less time on a given resource compared to specialists (18), and that generalists can suffer from information costs from having to track more information about their resource environment (19).

Another kind of cost that seems to be rarely considered is the cost of combining resources. We distinguish two types of combining costs. First, generalists occur in environments with shifting resource abundances and may need to switch among different resources. If different resources require different methods of exploitation, then changing among these methods can result in costs. These methods may be related to resource recognition, handling time, processing or detoxification (20, 21). We refer to these as resource-switching costs. Such costs have been observed in arctic charr, where the fish experience lower metabolic rates when fed a diet different from the one they were raised on compared to fish that did not have a dietary change (22).

A second kind of combining cost is when generalists try to feed on multiple resources at once and the specific techniques that work best for different resources are partly incompatible. It may be impossible to effectively deploy multiple optimal strategies at the same time. For example, synergistic interactions between defensive chemicals of different resources could occur such that the combined toxins are more detrimental (23). We refer to the reduction in foraging efficiency in the presence of multiple resources as resource-mixing costs. However, generalists could alternatively benefit from resource-mixing because of improved nutritional balance or through reduction in the amount of toxins ingested from any one resource or prey (24). These two kinds of combining costs aren’t the same as standard evolutionary trade-offs because a generalist may do just as well on every individual resource as a specialist but do more poorly when it tries to mix or switch among resources.

The costs and benefits associated with diet breadth evolution have been extensively studied in herbivorous insects. They are one of the most abundant and diverse eukaryotic life-forms and often possess highly specialized diets (25, 26). But the study of consumers with highly generalized diets provides an equally important perspective. Predators are often among the most general of consumers. By definition, predators are consumers that kill more than one victim in their lifetime (27), usually many more. They also tend to be much larger than their victims. Thus, to avoid starvation and minimize variance in energy intake, predators may need to consume many sub-optimal prey types when the most profitable prey is not abundant enough (28). Many macroscopic predators consume a formidable number of prey species, but few can match the diversity of prey consumed by some microbial protist predators. We call species with extreme diet breadth “super-generalists”. Predation by protists is a major factor accounting for bacterial mortality in the environment and as a consequence plays an important role in nutrient cycling (29–31). Predation by protists can determine the composition and properties of bacterial communities (32) and can be an important selective pressure for bacterial defenses such as biofilm formation, antibiotic production, and secretion systems (33, 34).

*Dictyostelium discoideum* is one such super-generalist protist predator. It is a social amoeba that lives in forest soils. It is a unicellular amoeba when bacterial prey are abundant and transitions to a non-feeding multicellular dispersal stage upon starvation. *D. discoideum* has been the subject of extensive research because of this fascinating multicellular stage (35, 36). Its feeding behavior has been less extensively studied, but it appears to qualify as a super-generalist predator. It can eat the majority of bacteria it is presented with (37–39). For example, one study tested 159 bacterial strains found in close association with fruiting bodies of *D. discoideum* from forest soil habitats and found that the amoebas were able to consume 77% of them (39).

The prey bacteria of *D. discoideum* are very diverse, ranging across at least four highly divergent bacterial phyla: Actinobacteria, Bacteroidetes, Firmicutes, and Proteobacteria (38). These shared a common ancestor about three billion years ago, far older than the divergence time of diverse insect prey (∼400 mya) that generalist invertebrates feed on (40). Bacteria also possess highly varied defensive mechanisms against microbial predators (33, 34). They can produce many kinds of secondary metabolite toxins to repel, disable, or kill their enemies. They can use secretion systems and effectors that can kill or allow for intracellular survival within protists. Some bacteria can swim away at high speeds to escape predators. Some can group together to form biofilms to prevent ingestion by predators.

Here we investigate the two kinds of costs of combining resources in *D. discoideum*. Amoebas occur in spatially and temporally variable communities of soil bacteria (41). Amoebas can encounter patchy bacterial distributions such that they switch from hunting one species of bacteria to another. We therefore tested for resource-switching costs by seeing if amoebas perform worse when switched to a new species of bacteria compared to controls that continued to grow on the same bacterium. Amoebas will often encounter mixed communities of prey, so we also looked at resource-mixing costs, testing if amoebas performed worse than expected in multi-species bacterial communities compared to their growth in single-species communities.

## Results

### Growth rate of *D. discoideum* amoebas varies on different species of bacteria

As a preliminary step we confirmed that *D. discoideum* is a generalist. We measured the growth rate of three *D. discoideum* clones on the commonly used lab food bacterium, *Klebsiella pneumoniae* and on 22 species of bacteria that had been collected as transient associates on *D. discoideum* fruiting bodies. These fruiting bodies emerged from soil and deer feces collected in the field, so the 22 bacterial species represent biologically relevant prey species for the amoebas. *D. discoideum* amoebas showed wide variation in their doubling times on different soil bacteria (Test bacterium: F_22,45_ = 23.88, p-value < 2.2×10^−16^). On each of the 23 bacteria, all three *D. discoideum* clones grew similarly (Figure 1), ruling out the possibility that the generalism of the species might be due to a mixture of individually specialized clones (42). The results also confirm that the amoebas are generalist feeders on prey bacteria that are likely to be encountered in nature.

**Figure 1.**
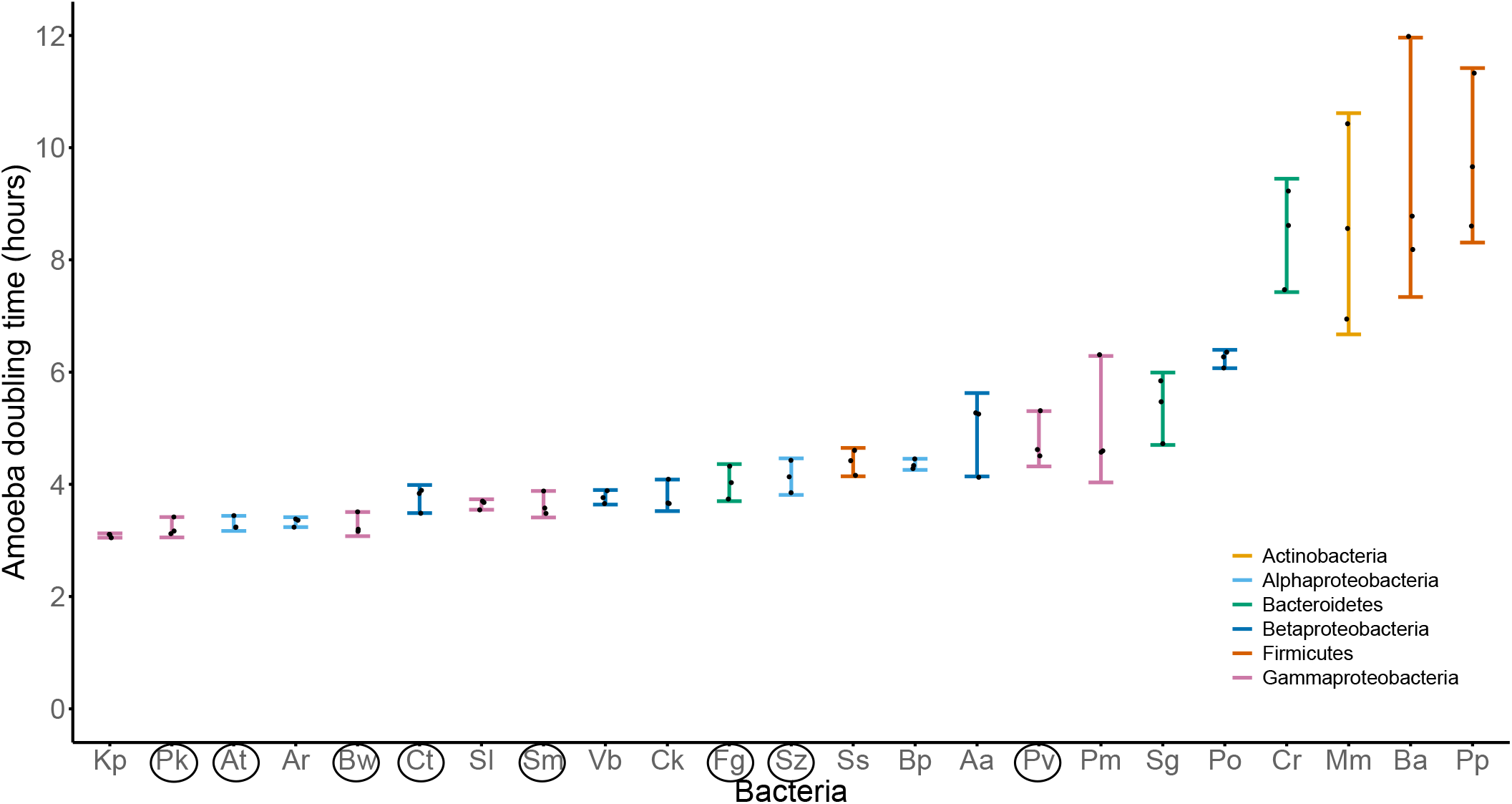
Doubling time of *D. discoideum* on *K. pneumoniae* and 22 species of soil bacteria closely associated with *D. discoideum*. Error bars are 95% C.I. The points represent the 3 *D. discoideum* strains. The circles represent the 8 species of bacteria used in other parts of this study. Bacteria species identities by Brock et al. 2018 based on closest partial 16S BLAST hit are listed in Table 1.

**Table 1:**
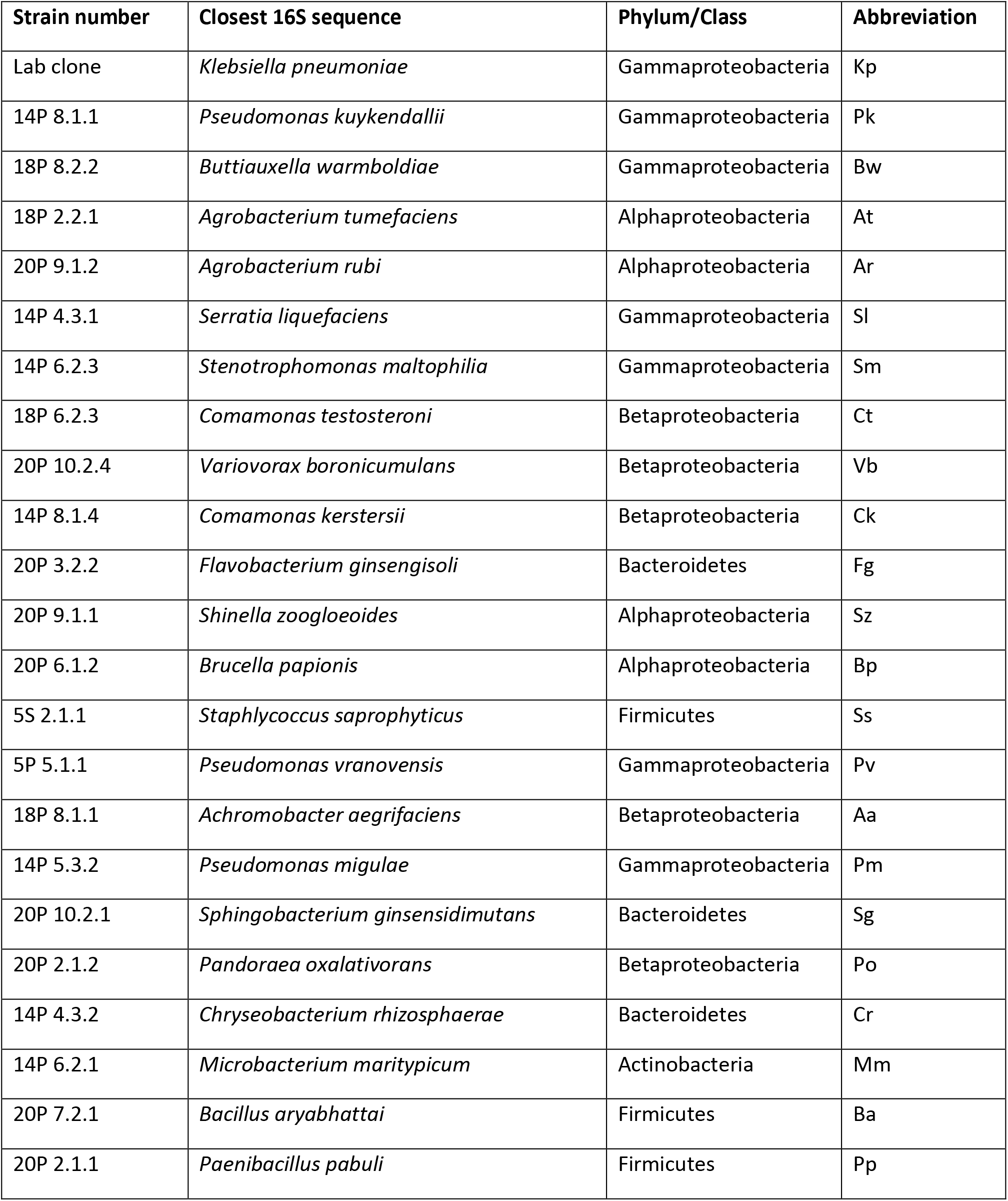
List of 23 species of bacteria used in *D. discoideum* growth rate experiments. * Species identification of bacterial isolates by Brock et al. 2018 based on closest partial 16S BLAST hit.

### *D. discoideum* amoebas experience resource-switching costs

We conducted resource-switching cost experiments to investigate how *D. discoideum* amoebas proliferate on a given bacterium when previously conditioned to a different bacterium (Figure 2). We found that there were fewer amoebas at 3 hours in switched treatments compared to controls that did not experience a prey switch (Figure 3, Treatment F_1,106_ = 27.621, p-value = 7.69×10^-7^; Effect size = −0.96, 95% CI = [−1.34, −0.575], df = 106; all effect sizes in this paper are the standardized measure Cohen’s d). Although only some bacterial species showed individually significant switching costs (Figure S1A), there was no interaction of treatment with prey species (Treatment x Test bacterium: F_1,106_= 0.838, p-value = 0.526) and the overall effect size was strong. Possible percentage change in amoeba numbers ranges from −24.04% to −8.03% (95% CI). We also found that conditioning time of amoebas (2 days or 5 days) had a similar effect on switching costs (Figure S1B, Conditioning time of 2 days: Treatment p-value < 0.001; Effect size = −1.094, 95% CI = [−1.63, −0.561], df =106, Conditioning time of 5 days: Treatment p-value = 0.0018; Effect size = −0.825, 95% CI = [−1.35, −0.301], df =106). This is inconsistent with switching costs being due to evolutionary changes during conditioning since evolutionary changes should be greater over 5 days than 2. The trend in the magnitude of effect sizes is also inconsistent with the evolutionary hypothesis because we would expect larger effect with longer conditioning time.

**Figure 2.**
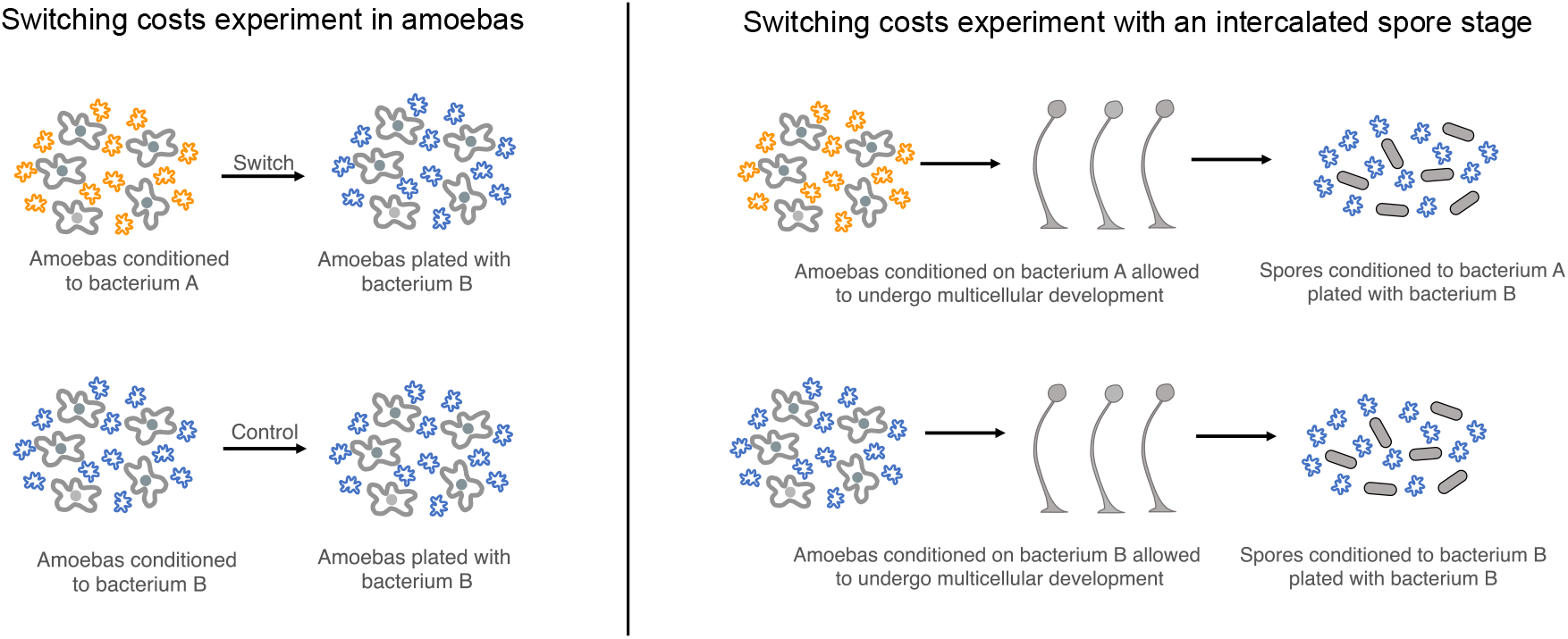
Schematic on the left shows resource-switching costs experiment in amoebas. Amoebas in switched treatment are conditioned on bacterium A and are switched to bacterium B for the experiment. The controls for this experiment are amoebas conditioned to bacterium B plated with a fresh culture of the same. We also varied conditioning time of amoebas on a given bacterium (2 days, 5 days). Schematic on the right outlines switching costs experiment on spores. We used the same basic switching design as described for the amoebas, except we let the amoebas undergo multicellular development on the conditioning bacterium. We then collected spores from these fruiting bodies and performed the experiment.

**Figure 3:**
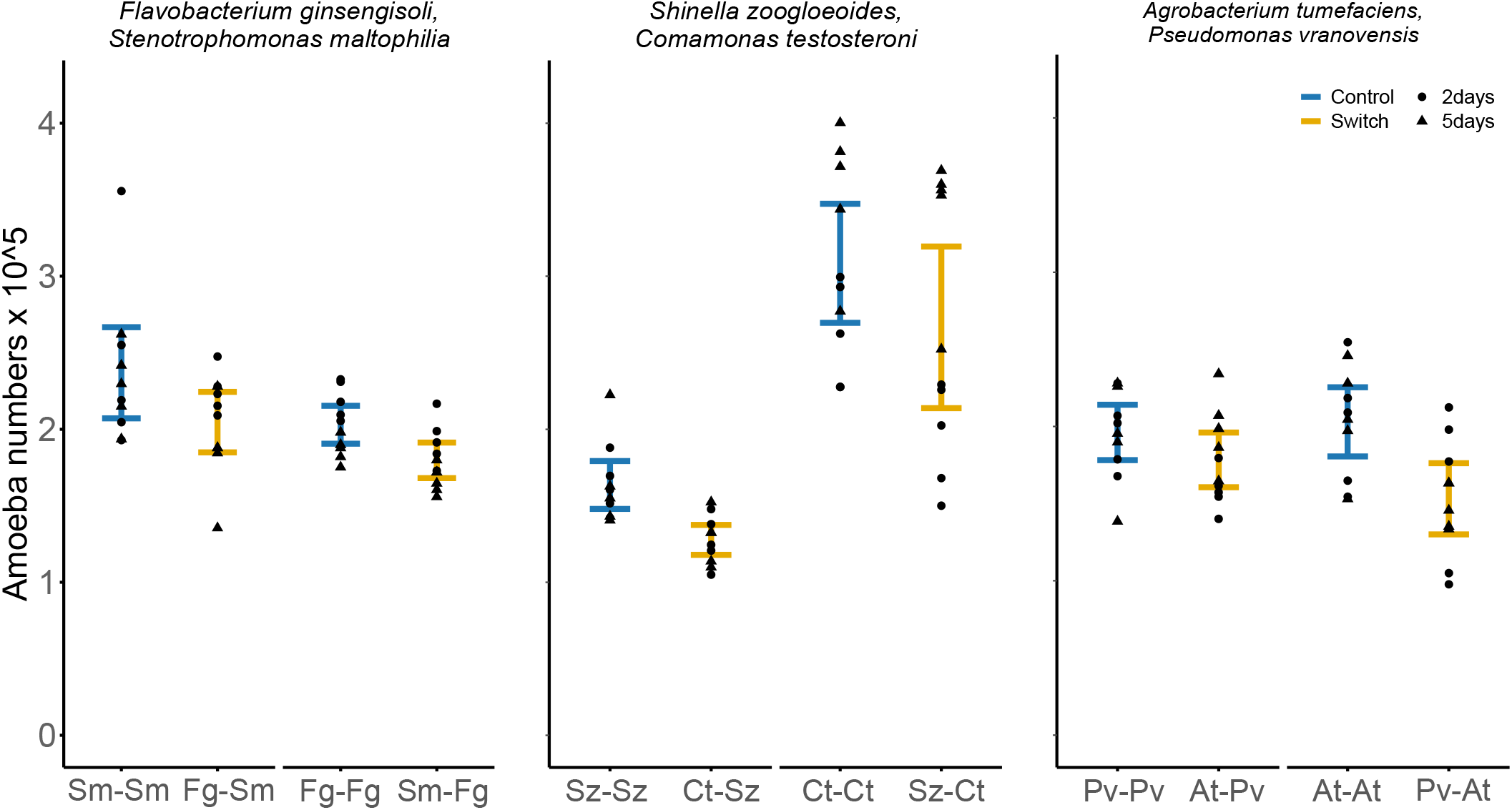
*D. discoideum* amoebas experience resource-switching costs: Mean amoeba numbers ± 95% C. I. of controls (blue) and switches (orange) after 3 hours of growth. The shapes represent the different conditioning times of amoebas (• - 2 days, ▴- 5 days). The points represent the five *D. discoideum* strains. There are fewer amoebas after three hours in switched treatments compared to controls. The two conditioning times are similar in the magnitude of switching costs. Interpreting x-axis labels: For example - Sm-Sm refers to amoebas grown on *S. maltophilia* moved to *S. maltophilia*, while Fg-Sm refers to amoebas grown on *F. ginsengisoli* moved to *S. maltophilia*.

### No evidence for resource-switching costs if amoebas undergo spore formation before the prey switch

If resource-switching costs are due to an evolved response in amoebas, then these genetic changes should be passed on through the spores. But if the costs are physiological, then undergoing the social cycle would be likely to erase the effects of food conditioning and therefore eliminate switching costs, because spores have very different gene expression profiles compared to amoebas (43, 44). We therefore tested if *D. discoideum* still show switching costs when they undergo the social cycle and spore formation before changing between prey bacteria. The switched and control spores resulted in a similar number of amoebas (Figure 4, Treatment: F_1,48_ = 0.263, p-value = 0.61; Effect size = −0.133, 95% CI = [−0.653,0.387], df = 48).

**Figure 4:**
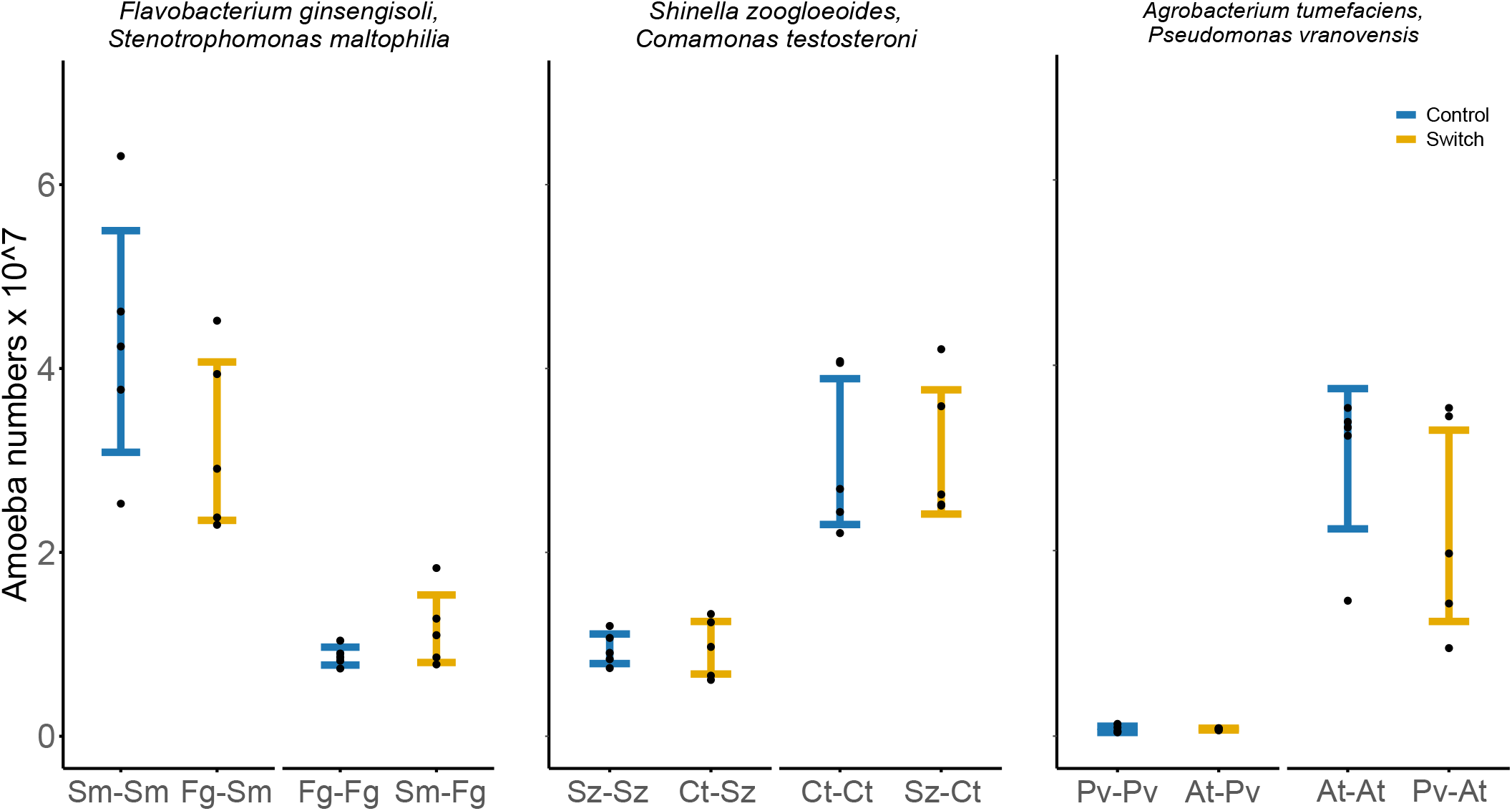
Switching costs disappear when spore formation precedes prey switch: the switched and control treatment have similar number of amoebas. Graph shows mean amoeba numbers ± 95% C.I. of controls and switches at 36 hours from plating. The points represent the five *D. discoideum* strains.

Possible percentage change in amoeba numbers ranges from −25.43% to 22.58%. Thus, we find no support for the hypothesis that switching costs are due to an evolved response, though the lower end of confidence interval of effect size does not rule out some cost.

### No evidence that resource-switching cost persists over the long term

If switching costs are not due to an evolved response, as supported by our previous results, then the switched amoebas should eventually recover their growth rates to the level of the controls that did not switch to new food bacteria. We conducted a time-course experiment to test this for all three pairs of bacteria and compared early (zero to six hours) and late (24 to 27 hours) growth rates of the switched and control treatments. As before (though tested here at 6 hours rather than 3 hours), the switched treatments have significantly lower early growth rates compared to controls (Figure 5A, Treatment F_1,53_ = 12.98, p-value = 6.9 x 10^-4^; Effect size =−0.931, 95% CI = (−1.48, −0.382), df= 53). Possible percentage change in growth rates ranges from - 27.47% to −3.75%. But there was no difference in late growth rate of switches and controls (Figure 5B, Treatment F_1,53_ = 0.014, p-value = 0.90; Effect size = −0.0311, 95% CI = (−0.549, 0.487), df= 53). Possible percentage change in growth rates ranges from −23.72% to 28.31%. Thus, though switched amoebas proliferate slower than controls during early growth, they appear to eventually catch up and perform similarly to controls, consistent with physiological adjustment to the new prey. Again, the confidence interval of effect size cannot rule out some cost at the late stage.

**Figure 5:**
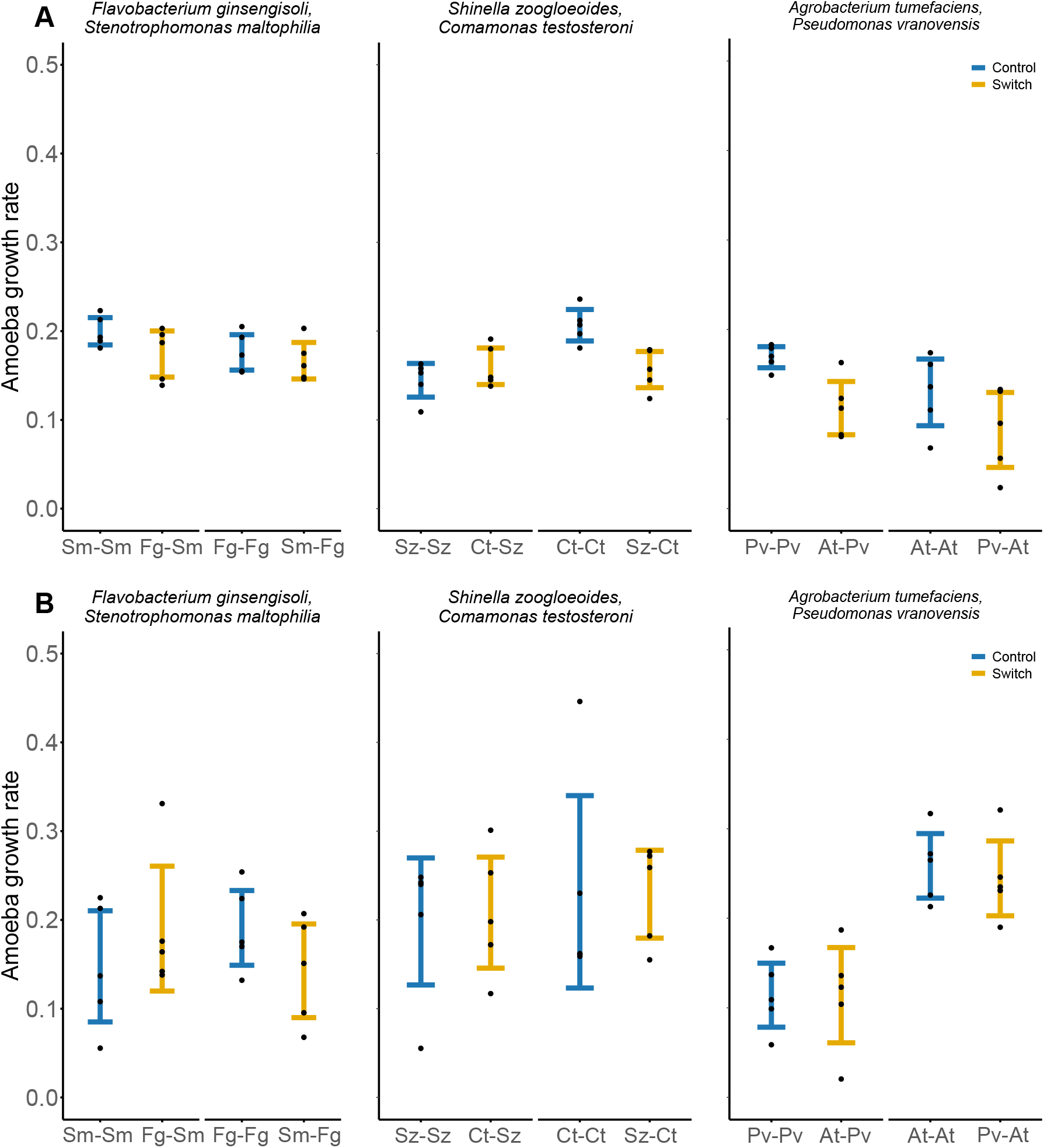
Switching costs occur during early growth period rather than later: **A)** Early growth rate estimates of controls and switches between 0 and 6 hours from set-up. Switched treatments have lower early growth rates compared to controls. **B)** Late growth rate estimates of controls and switches between 24 to 27 hours from set-up for. Graph shows mean growth rates ± 95% C.I. We found no difference in growth rates switches and controls at the later time points. The points represent 5 technical replicates of *D. discoideum* clone QS9.

### *D. discoideum* amoebas experience resource-mixing costs in some multi-prey communities

We tested if *D. discoideum* experiences costs when grown in multi-species prey communities compared to expectations from their growth in single-species communities. Consistent with resource-mixing costs, we observed significantly fewer amoebas than expected after growing in multi-species communities (Figure 6, Treatment F_1,59_ = 11.84, p-value = 0.001; Effect size = −0.811, 95% CI = [−1.31, −0.316], df = 59). Possible percentage change in observed and expected amoeba numbers ranges from −25.19% to −2.79%. But there seems to be variation in these mixing costs between the different bacterial communities (Treatment x Bacterial Community: F_5,59_ = 2.557 p-value = 0.036). Three of the prey communities cause significant costs as judged by non-overlap of effect size confidence intervals with zero, and none show a significant mixing benefit (Figure S2).

**Figure 6:**
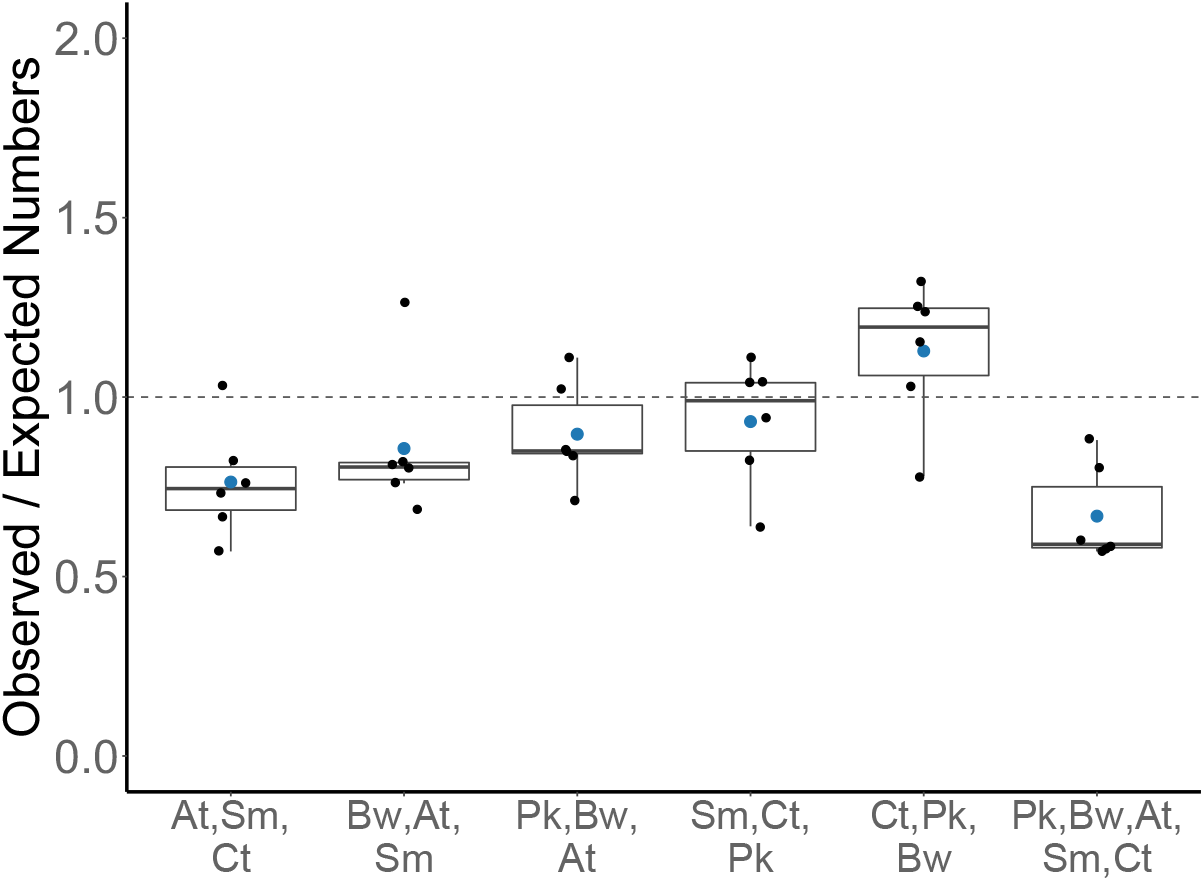
*D. discoideum* amoebas experience resource-mixing costs in some multi-species bacterial communites: Ratios of observed vs. expected amoeba numbers in different bacterial communities. A ratio lower than one indicates costs of resource mixing. The blue dot marks the mean. At- *Agrobacterium tumefaciens*, Bw- *Buttiauxella warmboldiae*, Ct - *Comamonas testosteroni*, Sm - *Stenotrophomonas maltophilia*, Pk - *Pseudomonas kuykendallii*.

## Discussion

Consumers lie on a spectrum between specialization on a few resources or generalism on a wide variety of resources. The balance between costs and benefits associated with these two strategies determines where the consumer falls in that spectrum and has important ecological and evolutionary consequences. But the costs of generalism are not well understood. We examined an understudied type of cost of diet generalism in the social amoeba, *D. discoideum*. As a preliminary step, we showed that the different strains of *D. discoideum* amoebas are true generalists that grow similarly on any given species of bacterium (Figure 1). Some have hypothesized that a generalist species could be a collection of individual specialists (23, 42), but our results show that this is not the case in *D. discoideum*.

Amoebas experience early resource-switching costs when moved from one prey bacterium to another, even after controlling for variation in bacterial edibility (Figures 3, 5A, S1A). The smallest costs consistent with the 95% confidence intervals were 8% and 4% and even these smallest effects seem biologically significant. We also did three experiments to test if these costs were physiological in nature or due to an evolved response. First, we found no evidence that the conditioning time (duration spent by amoebas on a given bacterium before the resource switch) affected the magnitude of switching costs. If the costs were due to evolution during the conditioning period, they should become stronger with longer conditioning time. Second, we found no evidence that amoebas experience switching costs if they undergo spore formation before prey switch (Figure 4). If this was due to an evolved response, then these effects would be genetically passed on through the spores. Third, we found that switching costs occur during early growth but are no longer apparent after a handful of additional generations (Figure 5), faster than they would change by selection. Our conclusions from these three experiments must be drawn from their failure to find significant costs, and the confidence intervals of each cannot exclude some costs. However, we have three different kinds of experiment, each one failing to support the hypothesis of evolutionary change and we have two separate experiments showing significant costs during the early stages where physiological effects were expected (Figures 3, 5A). Taken together these five experiments make a strong case that these costs are physiological in nature and not due to evolution.

Resource-switching costs have not been extensively studied and rarely enter into broader discussions of generalism, but there are other examples. Arctic charrs show reduced metabolic rate when switched among amphipods, bloodworms, and *Daphnia* (22). These costs might be related to irreversible changes in their digestive tracts during development which increase efficiency on the specific prey they were reared, but seem to be maladaptive on new prey. Some songbirds, such as American Robins and European Starlings, exhibit costs when switched between very different categories of foods in their natural diets, such as insects and fruit (45, 46). These switching costs can be reflected in utilization efficiencies, gut retention time, and body mass. Cabbage butterflies, *Pieris rapae*, take longer to extract nectar from a flower species if they are moved to it from a different flower (47). Switching between different flower species has a learning cost and may be related formation of a search-image for a specific flower. *Escherichia coli*, the model bacterium shows a distinct lag phase when shifting from one carbon resource to another (48). The lag phase may be a result of time required to switch on relevant metabolic genes or related to the ability of the cells to accurately detect the depletion of the primary carbon resource and presence of the secondary resource (49).

We also found that *D. discoideum* amoebas experience resource-mixing costs, where they proliferate less than expected in some communities with multiple species of prey bacteria. Our findings are in the opposite direction from the trend in other studies that have looked at the effect of mixing resources in generalists. Some studies on other protists that have looked at growth rate in multi-prey communities have found an increase in growth rates (50, 51).

Similarly, a study on marine amphipods found that fitness on mixed diets of algal species and animal matter was either improved or at least comparable to fitness on the best monospecific diet (52). A meta-analysis on the effect of diet mixing that included diverse consumers found that consumers grew better on mixed diets compared to the averages from single-species diet (53). However, if they considered only defended prey, then mixed diet was not different from the average of single-species diets. This might help explain our results. There could be negative synergistic interactions between defensive chemicals of different prey bacteria such that combined toxins are more detrimental to the amoebas. Another possible explanation of lower success on mixtures is that the different bacterial species compete and lower their overall numbers. However, we minimized this effect by not providing nutrients for bacterial growth. It is not clear why *D. discoideum* shows mixing costs when other taxa do not, but it is consistent with their switching costs.

It is interesting that we observe resource-mixing costs despite another factor that may obscure them. That factor is that amoebas may choose to eat the most profitable bacteria first. All but late measurements would then reflect the growth rate on the best bacterium and any mixing costs might therefore be obscured. Amoebas do show preferential attraction towards Gram-negative bacteria compared to Gram-positive bacteria (54). Thus, it would be interesting to test if *D. discoideum* amoebas can avoid some mixing costs by preferentially eating the most profitable prey bacteria first, but if they do, our data show that it is apparently not enough to fully overcome such costs.

These costs of combining resources are consistent with the idea that *D. discoideum* amoebas use partially different methods to hunt and process different prey. The costs probably involve changing gene expression. *D. discoideum* amoebas transcribe partially distinct sets of genes on Gram-positive (*Bacillus subtilis*, *Staphylococcus aureus*) and Gram-negative bacteria (*K. pneumoniae*, *Pseudomonas aeruginosa*) (55). The transcriptome of *D. discoideum* was also found to be highly species-specific when tested on 3 species of bacteria: *K. pneumoniae*, *B. subtlis*, and *Mycobacterium marinum* (56). Another study found that mutations that affect the ability of *D. discoideum* to grow on different bacteria were highly prey-specific (57). This suggests that there are different mechanisms for hunting or processing these distantly related bacteria.

Predation by amoebas is a complex process that can be divided into 4 broad steps: search, encounter, attack, and digestion (33). The costs of combining resources could arise in any of these steps. Eukaryotic phagocytes use G-protein coupled receptors (GPCRs) to detect and chase bacteria and use pattern recognition receptors to recognize and eliminate bacteria (58, 59). *D. discoideum* is equipped with 61 GPCRs (60). Some GPCRs are involved in the amoebas’ social cycle, but at least one GPCR, the folate receptor fAR1, has been implicated in both chasing and engulfing *Klebsiella* bacteria (61). Other GPCRs may play a role in other aspects of prey capture including chase and recognition. *D. discoideum* is also equipped with 22 genes that encode as many as 4 different types of lysozymes (62). Lysozyme genes play an important role in digesting bacteria. Knocking out some lysozyme genes generally reduces the ability of amoebas to feed on all tested bacteria, while deletion of other lysozyme genes reduces growth only on specific gram-negative bacteria (63). Future research on the mechanisms underlying these costs would be valuable.

Why can’t amoebas evolve one predation technique that works optimally on all prey bacteria to eliminate the costs of diet generalism? After all, amoebas are a few hundred times larger than the average bacterium by mass (64). They should be able to overwhelm most bacteria with their size advantage. But bacteria possess many mechanisms to resist their eukaryotic predators (33, 34, 65, 66). Morphological adaptations such as formation of microcolonies, biofilms, and filamentation can prevent attack or ingestion by predators. Bacteria can modify their membrane properties by changing the lipopolysaccharides on the outer membrane, secreting an S-layer and sporulating among others that can help them avoid recognition, ingestion, and digestion by predators. They can also produce a huge array of secondary metabolites that can deter predation by protists (67, 68). Thus, each bacterial species may possess a unique combination of defenses that is unlikely to yield to a common predation strategy. It would be interesting to test how many different predation strategies *D. discoideum* can employ. *D. discoideum* may treat some groups of bacteria similarly and some as different.

We suspect that environmental heterogeneity plays a large role in the maintenance of a generalist strategy in *D. discoideum*. Environmental heterogeneity, especially temporal variation favors generalists (69). The scale of variation can also influence the nature of generalism (70). Coarse-grained environments may select for early developmental plasticity, but for a fixed phenotype later in life. Fine-grained environments select for versatile generalists that are capable of reversing their phenotypic response.

How much environmental heterogeneity amoebas experience in nature is not fully known. Soil bacterial communities are certainly spatially and temporally variable, such that no one bacterium may be consistently sufficiently abundant (41). But the scale is important. On a microscale, soil is made up of small aggregated particles that are connected via a network of air- and water-filled pores (71). The three-dimensional nature of these particles increases soil surface area such that as little of 10^-6^ percent of soil surfaces may contain bacteria (72). These particles generally contain bacterial patches of a limited number of clonal cells and these patches can be separated by distances that are large on a microbial scale (73). Thus, it is possible that amoebas are likely to experience switching costs as they move between bacterial patches. Amoebas can also travel considerable distances during their social cycle through slug migration and especially through the dispersal of spores by animal vectors (74). This can introduce them to new soil environments with different bacterial communities. Thus, amoebas are in a selective environment that probably favors generalists.

Resource-combining costs may be less relevant to consumers that can find one big resource unit to spend their whole life on, such as phytophagous insects or parasitoid wasps. However, the generalist parasitoid wasp, *Venturia canescens* shows a cost of resource switching across generations, perhaps due to epigenetic or maternal effects (75). Consumers that encounter diverse and defended prey species in their lifetimes are more likely to be plagued by these costs. One example is wolf spiders that experience costs when fed a mixed diet of toxic prey (76).

Resource-combining costs are different from standard trade-offs because here generalists can handle each resource well, but they experience costs when combining different resources. These costs, like trade-offs, could eventually lead to resource specialization. We have shown that resource-combining costs are prominent in a resource generalist but it would be interesting to also study resource specialists. Are they sometimes constrained in their diets less by being unable to feed on a more general diet, but by the inability to readily switch to another suitable resource? Are they kept from having a broader diet by resource-mixing costs? We suggest that resource-combining costs deserve greater consideration than they have received in the debate on the evolution of generalism and specialism.

## Materials and Methods

### Growth rate of *D. discoideum* on different species of bacteria isolated from forest soil environments

We first measured the growth rate of wild clones of *D. discoideum* on *Klebsiella pneumoniae,* the lab food bacterium, and on 22 species of bacteria found in close-association with *D. discoideum* isolated by Brock et al. 2018 from forest soil environments (Table 1). These bacteria were isolated from fruiting bodies of *D. discoideum* that developed from field-collected samples of soil and deer feces from Mountain Lake Biological Station, Virginia in 2014. All *D. discoideum* strains used in this paper were also isolated from Mountain Lake Biological Station, Virginia. The bacterial species identification is based on closest partial 16S BLAST hit by Brock et al. 2018. In this paper, we refer to the bacteria using names from these closest 16s BLAST hits. We performed this experiment on 3 *D. discoideum* strains: QS1, QS6, and QS9. We plated 100,000 amoebas with 200μl of 10 OD_600_ bacterial suspension on starving agar plates. We estimated amoeba numbers at 20 hours and calculated doubling times as:

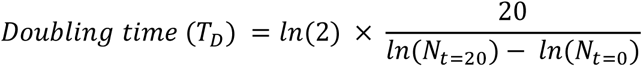

This assay also helped us identify reasonably edible bacteria for other experiments.

### Resource-switching cost assay in *D. discoideum* amoebas

To test if resource-switching costs occur when *D. discoideum* amoebas are switched from Bacterium A to Bacterium B, we first conditioned separate populations of the amoebas to each species of bacterium (Figure 2, details below for three separate experiments). The switched treatment for these experiments used amoebas conditioned to Bacterium A and replated with Bacterium B. We plated 100,000 amoebas conditioned to Bacterium A with 200μl 20 OD_600_ suspension of Bacterium B on SM/20 plates and then measured amoeba numbers after 3 hours. The controls for this experiment were identical except that amoebas conditioned to Bacterium B were replated with Bacterium B. If we observe fewer amoebas in switched plates compared to control plates, then *D. discoideum* amoebas experience resource-switching costs. We measured the costs of switching between 3 pairs of bacteria with 2 reciprocal switches for each pair. We replicated this experiment with 5 *D. discoideum* strains: QS6, QS9, QS14, QS17, QS160.

We did three sets of experiments to rule out the possibility that these costs are instead due to an evolved response in the amoebas during the conditioning period to either bacterium. Switched amoebas could experience poor growth on a Bacterium B because of trade-offs associated with new evolutionary adaptation to the Bacterium A during the conditioning period. Control amoebas may also experience better growth rates if they have evolved and adapted to Bacterium B during the conditioning period. We ruled out this complication in three ways.

### Effect of conditioning length of amoebas on resource-switching costs

First, we tested if the length of the conditioning period of amoebas on the bacterium affected resource-switching costs. If costs are due to either adaptation in switched amoebas that results in evolutionary trade-offs, or to direct improvement in growth rate because of adaptation in control amoebas, we expect stronger costs with longer conditioning (i.e. evolving) time. Thus, we conditioned amoebas on a given bacterium for a) 2 days and b) 5 days before measuring switching costs. In the 2-day treatment, we conditioned the amoebas for 40 hours by plating 200,000 spores of *D. discoideum* with 200ul of 1.5 OD_600_ bacterial suspension on SM/5 plates. After ∼40 hours, we washed off the plates with 10ml of ice cold KK2 to collect the amoebae-bacteria suspension. Next, we centrifuged the suspension for 3 minutes at 300g at 10°C to spin down the amoebas and discarded the supernatant containing the bacteria. We resuspended the amoeba pellet in ice-cold KK2 and washed it off two more times until all the bacteria were discarded. Finally, we suspended the amoeba pellet in 500μl – 1000μl of ice-cold KK2. We conditioned the amoebas for the 5-day experiment similarly, but collected amoebas from the resulting vegetative front after 5 days with a sterile loop and resuspending in 600ul ice-cold KK2. We performed washing centrifuging steps as described above to thoroughly wash the amoebas off the conditioning bacterium. A conditioning time of 2 days translates to 10−14 amoeba divisions and 5 days to 25-35 amoeba divisions on good prey. If resource-switching costs are because of an evolutionary response, then we expect costs to be stronger for the 5-day conditioning period than the 2-day period.

### Resource-switching costs assay with *D. discoideum* spores

To further distinguish between a temporary switching cost and an evolved response, we tested whether allowing amoebas to go through the social cycle and subsequent sporulation before prey bacteria switches erases switching costs (Figure 2). If costs are because of an evolved response in amoebas, then these changes should still be evident after the social stage. But if the costs are because of conditioning of the amoebas that results in temporary mismatch in transcriptional tools on the new bacterium, then undergoing the social cycle should erase most switching costs. This is because *D. discoideum* experiences a huge turn-over in gene expression when transitioning from a unicellular lifestyle into the multicellular cycle (Parikh et al. 2010, Rosengarten et al. 2015). The abundance of almost every transcribed mRNA changes at least 2-fold throughout development starting from vegetative amoebas to multicellular fruiting bodies (43). Therefore, any transcriptional conditioning of amoebas towards a bacterium should be largely erased by development. The switched and control spores are at the same transcriptional start line of dormancy.

We used the same design as the switching experiment for amoebas described above but with sporulation at the end of the conditioning phase (∼7 days). *D. discoideum* amoebas feed on bacteria for the first 2-4 days of the conditioning phase, and then transition to the social cycle once the bacteria are depleted. We then plated 100,000 spores from the resulting fruiting bodies with bacterial suspensions on SM/20 (Figure 2). Since the spores require some time to germinate into amoebas and start feeding on the bacteria, we counted the total number of amoebas on these plates after 36 hours.

### Time course data of resource-switching costs in *D. discoideum* amoebas

As a third check on whether what we see is an evolved or a conditioned response, we conducted a time series study of the amoebas on all three pairs of bacteria to check if the growth rates of the switched treatments quickly catch up with control growth rates, which would indicate a physiological lag rather than an evolutionary change. We tracked the number of amoebas in switched and control plates after 6, 24 and 27 hours from plating. We calculated early growth rate between 0h and 6h time points and calculated late growth rate between 24h and 27h time points. We assumed exponential growth and used this formula to calculate growth rate between time points t1 and t2, where N_t_ stands for number of amoebas at time t:

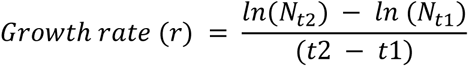

If switching costs are not an evolved response, then switches and controls should not differ in their late growth rates. We performed this on SM/5 plates with a starting number of 100,000 amoebas with 200μl of 1.5 OD_600_ suspension of the test bacterium on the *D. discoideum* clone QS9. We replicated this experiment five times.

### Resource-mixing costs in *D. discoideum* amoebas

Next, we determined whether *D. discoideum* amoebas experience resource-mixing costs when feeding on multiple species of bacteria. For this experiment, we chose 5 prey bacteria that were generally highly edible from our growth rate assay of amoebas on 23 species of bacteria. We generated 5 single-species communities, 6 three-species communities and 1 five-species community from the 5 prey bacteria. We made 10 OD_600_ suspensions of each bacterial species and mixed these suspensions in equal proportions to make a given multi-species suspension. We used AX4 amoebas that were grown axenically in bacteria-free cultures to preclude any effect of prior conditioning of amoebas to a given species of bacterium on the assay. We plated 100,000 amoebas on starving agar plates with 200μl of bacterial suspensions and measured amoeba numbers in all the different treatments after 20 hours. We replicated this experiment 6 times.

We expect bacterial growth and consequently competition to be minimal because we performed this experiment on starving agar. To test if amoebas experience resource-mixing costs, we first calculated the expected number of amoebas in multi-prey treatments with data from our single-prey treatments. For example, the expected number of amoebas in multi-prey treatment containing *A. tumefaciens*, *S. maltophilia* and *C. testosteroni* is the average of observed numbers of amoebas in single-prey treatment of those bacteria. We then compared these expected numbers to our observed number of amoebas in these multi-species treatments to infer costs.

### Statistics

We performed all statistical analysis in R (version 4.2.1) (77). We used general linear models (with log link functions for count data) to analyze our data after testing for normality of residuals using the Shapiro-Wilk’s test and examining Q-Q plots. We used the “*emmeans*” package to calculate estimated effect sizes (Cohen’s d, a standardized measure) and 95% confidence intervals for relevant contrasts (78). We calculated the largest and the smallest percentage difference between treatments from the 95% C.I range of the estimated marginal means. To test if *D. discoideum* amoebas experience varying doubling times on different species of bacteria, we built a linear model with amoeba doubling times as the response variable and the Test bacterium (23 species of bacteria listed in Table 1) as the fixed factor. We define test bacterium as the bacterium on which we measured amoeba growth rates. To test if *D. discoideum* amoebas experience a resource-switching cost on a given bacterium when previously grown on a different bacterium, we built a linear model with log-transformed amoeba numbers after 3 hours of growth as the response variable and Treatment (Control, Switch), Test bacterium (At, Pv, Fg, Sm, Sz, Ct), and Conditioning length (2days, 5days) as fixed factors. We also included interaction effects for Treatment x Test bacterium, Treatment x Conditioning length. For the similar experiment that includes a spore stage after conditioning, we used a similar linear model with log transformed amoeba numbers after 36 hours as the response variable and Treatment and Test bacterium as the fixed factors. We included interaction effects for Treatment x Test bacterium in this model.

We performed the following statistical tests on time-course data collected on switches. To test if there are switching costs during early proliferation we used a linear model with early growth rate calculated between 0 and 6 hours as the response variable and Treatment (Control, Switch) and Test bacterium (At, Pv, Fg, Sm, Sz, Ct) as fixed factor. To test if resource-switching costs persist during late proliferation, we used a similar linear model with late growth rate calculated between 24 and 27 hours as the response variable.

To test if *D. discoideum* amoebas experience resource-mixing costs, we tested if amoebas performed worse than expected in multi-species prey communities. We built a linear model with log transformed amoeba numbers at 20 hours as the response variable, Treatment (categorical variable for whether an observation is expected or observed) and Bacterial Community (6 Communities) and Experimenter (2 Experimenters) as fixed factors. We included an interaction effect between Treatment x Bacterial Community. ANOVA tables for all models are included in the supplement.

## Acknowledgments

This material is based upon work supported by the National Science Foundation under grants IOS 16-56756, DEB 17-53743, and DEB 2237266. We thank Trey Scott for feedback on statistical analysis. We thank Margaret Steele and Calum Stephenson for comments on the manuscript.

We also thank the Queller-Strassmann lab for their helpful insights on experimental design.

## Author Contributions

P.M.S., J.E.S., and D.C.Q. designed research. P.M.S., D.A.B., and R.I.M. performed the experiments. P.M.S., J.E.S., and D.C.Q. analyzed data. P.M.S., J.E.S., and D.C.Q. wrote the paper.

## Classification

Major - Biological Sciences, Minor-Ecology

---

**Figure S1:**
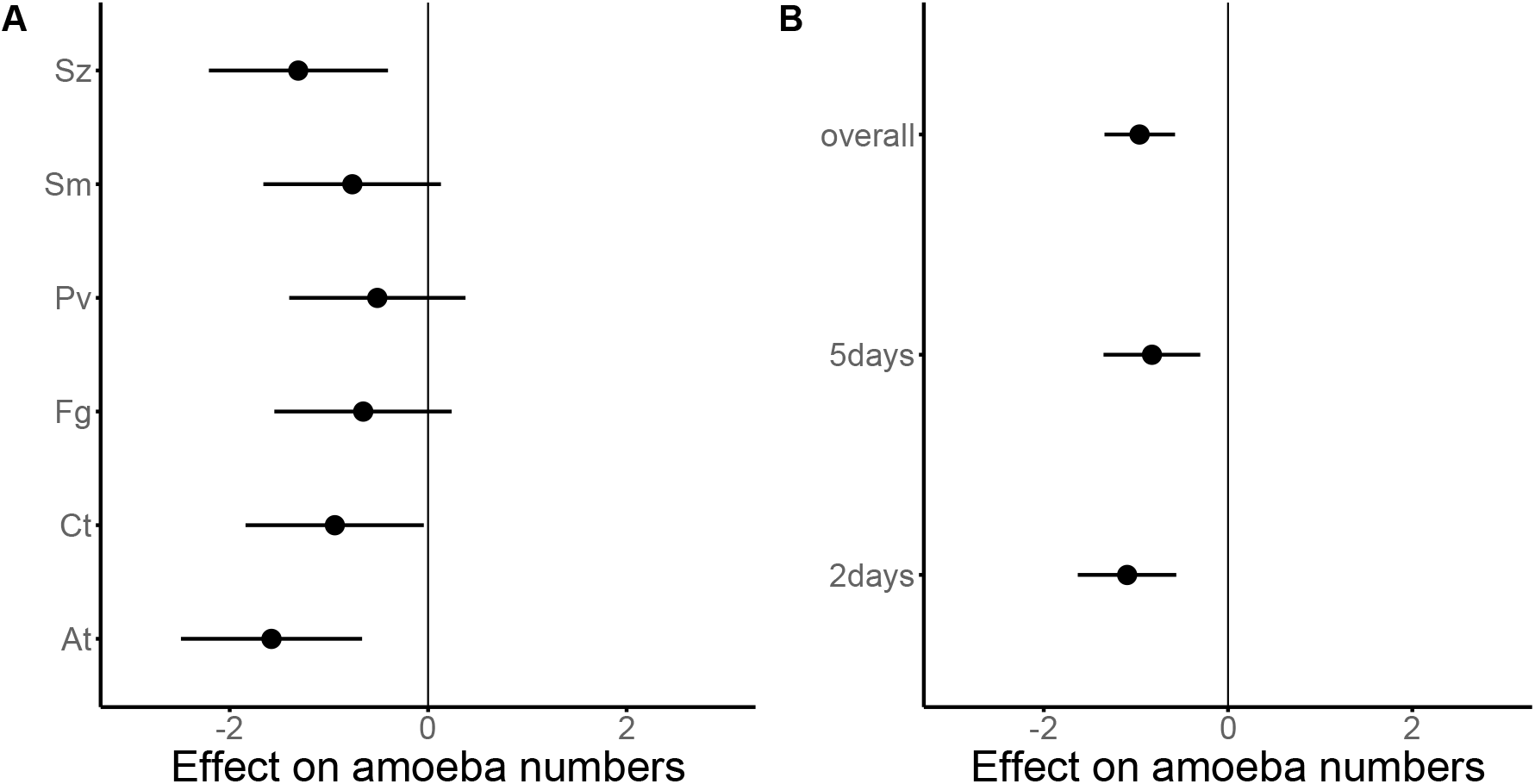
Estimated effect size (Cohen’s d) ± 95% C.I. of amoeba numbers of the switched treatment minus the control treatment. **A)** shows that switched treatment in all test bacterium generally had a roughly similar negative effects on amoeba numbers. Plot **B)** shows that switched treatments of the different conditioning time had a similar negative effect on amoeba number compared to controls.

**Figure S2:**
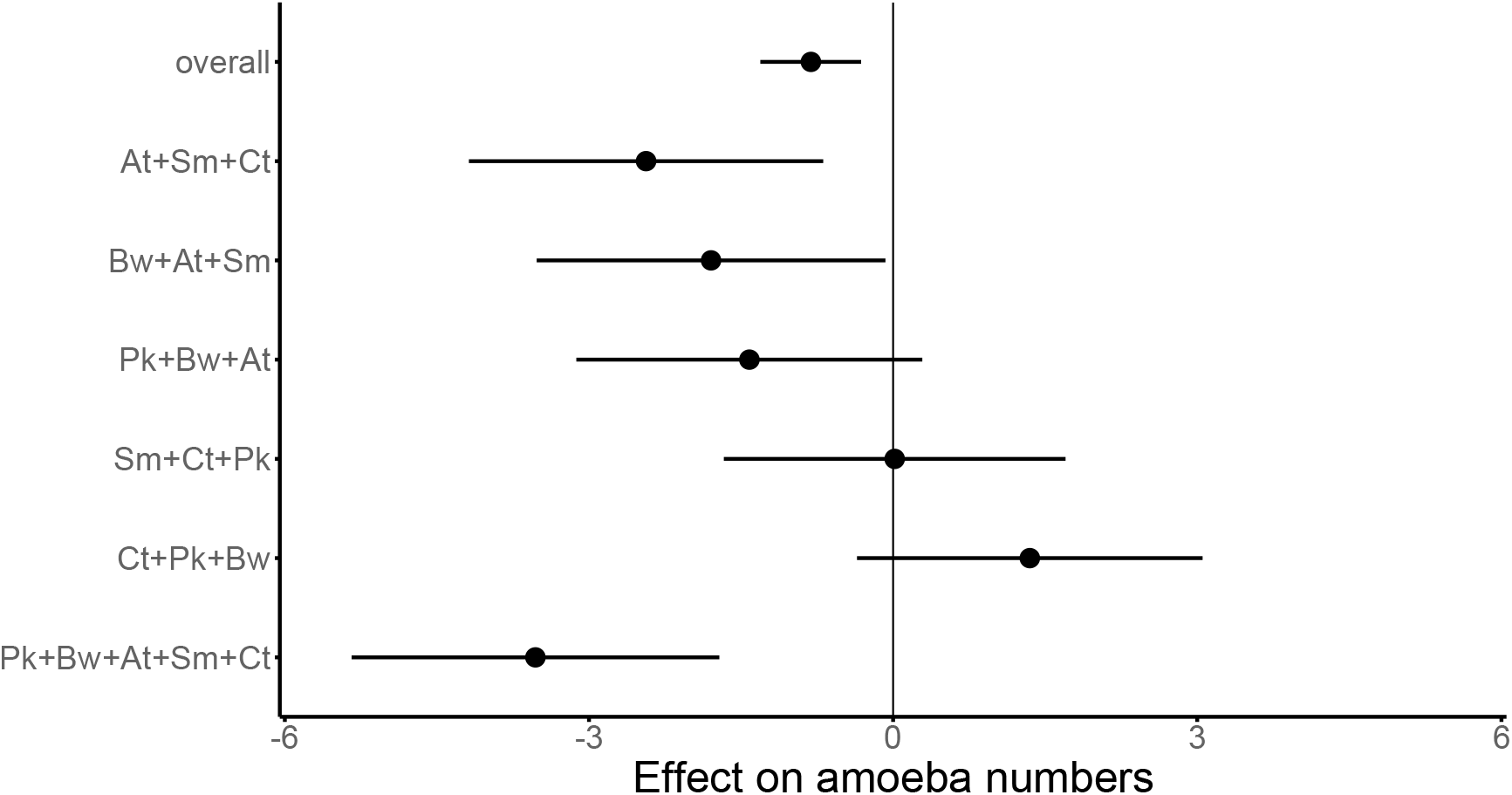
Estimated effect size (Cohen’s d) ± 95% C.I. of observed – expected amoeba numbers in resource-mixing experiment averaged across observers. We see support for mixing costs overall and expected trend in four out of the six bacterial communities.

**Table S1:** ANOVA of amoeba growth rate assay. Formula: anova(lm (log(amoeba_number) ~ Test_bacterium))

|  | Df | Sum sq | Mean Sq | F_value | Pr(>F) |
| --- | --- | --- | --- | --- | --- |
| Test_bacterium | 22 | 299.346 | 13.3794 | 24.949 | 2.20E-16 |
| Residuals | 46 | 24.668 | 0.5363 |  |  |

**Table S2:** ANOVA of amoeba switching assay. Formula: anova(lm(log(amoeba_number) ~ Treatment x Test_bacterium + Treatment × Conditioning_length)

|  | Df | Sum sq | Mean Sq | F_value | Pr(>F) |
| --- | --- | --- | --- | --- | --- |
| Treatment | 1 | 0.965 | 0.9648 | 27.621 | 7.69E-07 |
| Test_bacterium | 5 | 4.958 | 0.9917 | 28.392 | 2.00E-16 |
| Conditioning_length | 1 | 0.057 | 0.0567 | 1.623 | 0.205 |
| Treatment x Test_bacterium | 5 | 0.146 | 0.0293 | 0.838 | 0.526 |
| Treatment x Conditioning_length | 1 | 0.019 | 0.019 | 0.543 | 0.463 |
| Residuals | 106 | 3.702 | 0.0349 |  |  |

**Table S3:** ANOVA of spore switching assay. Formula: anova(lm(log(amoeba_number) ~ Treatment × Test_bacterium + Treatment × Conditioning_length))

|  | Df | Sum sq | Mean Sq | F_value | Pr(>F) |
| --- | --- | --- | --- | --- | --- |
| Treatment | 1 | 0.03 | 0.03 | 0.264 | 0.61 |
| Test_bacterium | 5 | 106.78 | 21.355 | 186.402 | <2e-16 |
| Treatment x Test_bacterium | 5 | 0.68 | 0.136 | 1.191 | 0.328 |
| Residuals | 48 | 5.5 | 0.115 |  |  |

**Table S4:** ANOVA of early growth rate data. Formula: anova(lm(log(amoeba_number) ~ Treatment + Test_bacterium))

|  | Df | Sum sq | Mean Sq | F_value | Pr(>F) |
| --- | --- | --- | --- | --- | --- |
| Treatment | 1 | 0.012 | 0.012 | 12.988 | 0.0006923 |
| Test_bacterium | 5 | 0.043 | 0.009 | 9.561 | 1.52E-06 |
| Residuals | 53 | 0.048 | 0.001 |  |  |

**Table S5:**
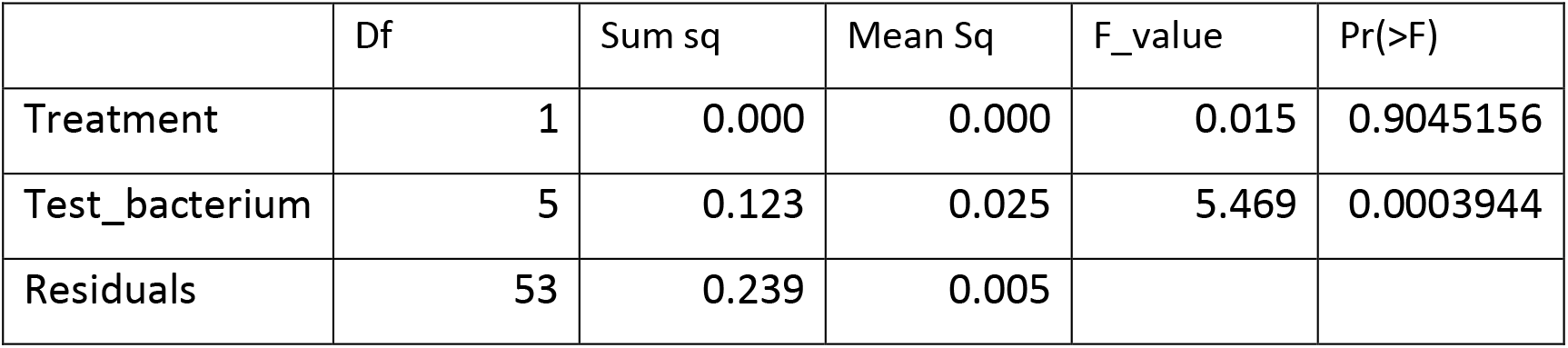
ANOVA of late growth rate data. Formula: anova(lm(log(amoeba_number) ~ Treatment + Test_bacterium))

**Table S6:** ANOVA of resource-mixing data. Formula: anova(lm(log(amoeba_number) ~ Treatment *Community + Experimenter))

|  | Df | SumSq | MeanSq | F-value | Pr(>F) |
| --- | --- | --- | --- | --- | --- |
| Treatment | 1 | 0.4564 | 0.4564 | 11.8394 | 0.001071 |
| Community | 5 | 1.6847 | 0.3369 | 8.74 | 3.01E-06 |
| Experimenter | 1 | 24.3675 | 24.3675 | 632.0609 | 2.20E-16 |
| Treatment x<br>Community | 5 | 0.4929 | 0.0986 | 2.5571 | 0.036767 |
| Residuals | 59 | 2.2746 | 0.0386 |  |  |

